# Fast genome-wide functional annotation through orthology assignment by eggNOG-mapper

**DOI:** 10.1101/076331

**Authors:** Jaime Huerta-Cepas, Kristoffer Forslund, Damian Szklarczyk, Lars Juhl Jensen, Christian von Mering, Peer Bork

## Abstract

Orthology assignment is ideally suited for functional inference. However, because predicting orthology is computationally intensive at large scale, and most pipelines relatively inaccessible, less precise homology-based functional transfer is still the default for (meta-)genome annotation. We therefore developed eggNOG-mapper, a tool for functional annotation of large sets of sequences based on fast orthology assignments using precomputed clusters and phylogenies from eggNOG. To validate our method, we benchmarked Gene Ontology predictions against two widely used homology-based approaches: BLAST and InterProScan. Compared to BLAST, eggNOG-mapper reduced by 7% the rate of false positive assignments, and increased by 19% the ratio of curated terms recovered over all terms assigned per protein. Compared to InterProScan, eggNOG-mapper achieved similar proteome coverage and precision, while predicting on average 32 more terms per protein and increasing by 26% the rate of curated terms recovered over total term assignments per protein. Through strict orthology assignments, eggNOG-mapper further renders more specific annotations than possible from domain similarity only (e.g. predicting gene family names). eggNOG-mapper runs ~15x than BLAST and at least 2.5x faster than InterProScan. The tool is available standalone or as an online service at http://eggnog-mapper.embl.de.

## Introduction

The identification of orthologous genes, originating from speciation rather than duplication events (Fitch 1970), is a long-standing evolutionary problem with deep implications for the functional characterization of novel genes. The ‘Ortholog Conjecture’ states that ancestral functions are more likely to be retained between orthologous genes than between paralogs (Tatusov et al. 1997). Therefore, information gained on the role of a gene in a model organism is potentially transferrable to its orthologs in less experimentally tractable species. While this motivation remains central (Gabaldón and Koonin 2013), its application is frequently left up to users (e.g. genome annotators) in the form of *ad hoc* scripted solutions, often based on more general homology searches rather than orthology assignments. Most tools in use for functional annotation of newly sequenced genomes apply BLAST (Blast2GO (Götz et al. 2008); RAST (Overbeek et al. 2014)) or HMMER searches (Finn et al. 2014; Jones et al. 2014) to transfer functional terms from homologous sequences.

Building on recent improvements made to the eggNOG orthology resource (Huerta-Cepas et al. 2016), we have created eggNOG-mapper, an application intended for fast functional annotation of novel sequences using precomputed sequence profiles and orthology assignments. The tool is designed for the annotation of large collections of novel sequences, typically targeting translated gene-coding regions from (meta-)genome and transcriptome data.

## New Approaches

The annotation algorithms in eggNOG-mapper are implemented as follows:

1. Sequence Mapping (Figure 1A). For each query sequence, HMMER 3 (Eddy 2011) is first used to search for significant matches in the precomputed collection of Hidden Markov Models (HMM) available from the eggNOG database (Huerta-Cepas et al. 2016). HMM matches, each associated to a functionally annotated eggNOG Orthologous Group (OG), provide a first (more general) layer of functional annotation. Next, each query protein is searched against the set of eggNOG proteins represented by the best matching HMM using the *phmmer* tool. Finally, the best matching sequence for each query is stored as the query’s seed ortholog and usedtoretrieveother orthologs (see step 2 below). At present, eggNOG HMM collection comprises sequence profiles of 1,911,745 Orthologous Groups (OGs), spanning 1678 bacteria, 115 archaea, 238 eukaryotes and 352 viruses. 104 sub-databases are available that allow restricting searches to narrower taxonomic groups, thereby speeding up computations and enforcing annotations to be exclusively transferred from orthologs in a particular set of species. Alternatively, a faster mappingapproachcanbe selected that uses DIAMOND (Buchfink et al. 2015) to search for the best seed ortholog of each query directly among all eggNOG proteins. This option is considerably faster than the HMM approach, but should be considered less sensitive (e.g. HMMER sensitivity is comparable to PSI-BLAST (Söding 2005) whereas DIAMOND is comparable to BLAST). Although the use of DIAMOND had little impact in the re-annotation of model organisms (see additional benchmarks in Supplementary Material), novel sequencescoming fromorganisms without close representatives in the eggNOG taxonomic scope may indeed be overlooked. HMMER searches, however, would still allow for functional annotation of distant homologs via HMM matches.
2. **Orthology assignment** (Figure 1B). For each query, the best matching sequence, which points to a protein in eggNOG, is used to retrieve a list of fine-grained orthology assignments from a database of pre-analysed eggNOG phylogenetic trees (i.e. excluding evident (in-)paralogs as described in (Huerta-Cepas et al. 2007)). Additional filters such as bit-score or E-value thresholds can be used at this step in order to avoid inferring functional data for query sequences without sufficient homology to any protein in the eggNOG database.
3. **Functional Annotation.** All functional descriptors available for the retrieved orthologs are transferred to the corresponding query proteins. By default, functional transfers are automatically restricted to the taxonomically closest orthologs of each query, reducing the risk of false assignments from too distant species (Figure 1C).Thisparameterisautomaticallyadjustedfor every sequence, without the need of prefixing any taxonomic filter and allowing each query to be annotated using the most suitable taxonomic source. Finally, although all orthologs are considered by default, users can choose to restrict annotations to those based on one-to-one orthology assignments only (Figure 1D, left), thus increasing the reliability of functional transfers at the cost of lower annotation coverage (see Supplementary Materials). Functional descriptors are based on the most recent eggNOG build, and currently include curated GO terms (Gene Ontology Consortium 2015), KEGG pathways (Kanehisa et al. 2014) and COG functional categories (Galperin et al. 2015). Moreover, taking advantage of the fine-grained orthology assignments, gene family names are predicted for each query.

**FIGURE 1.**
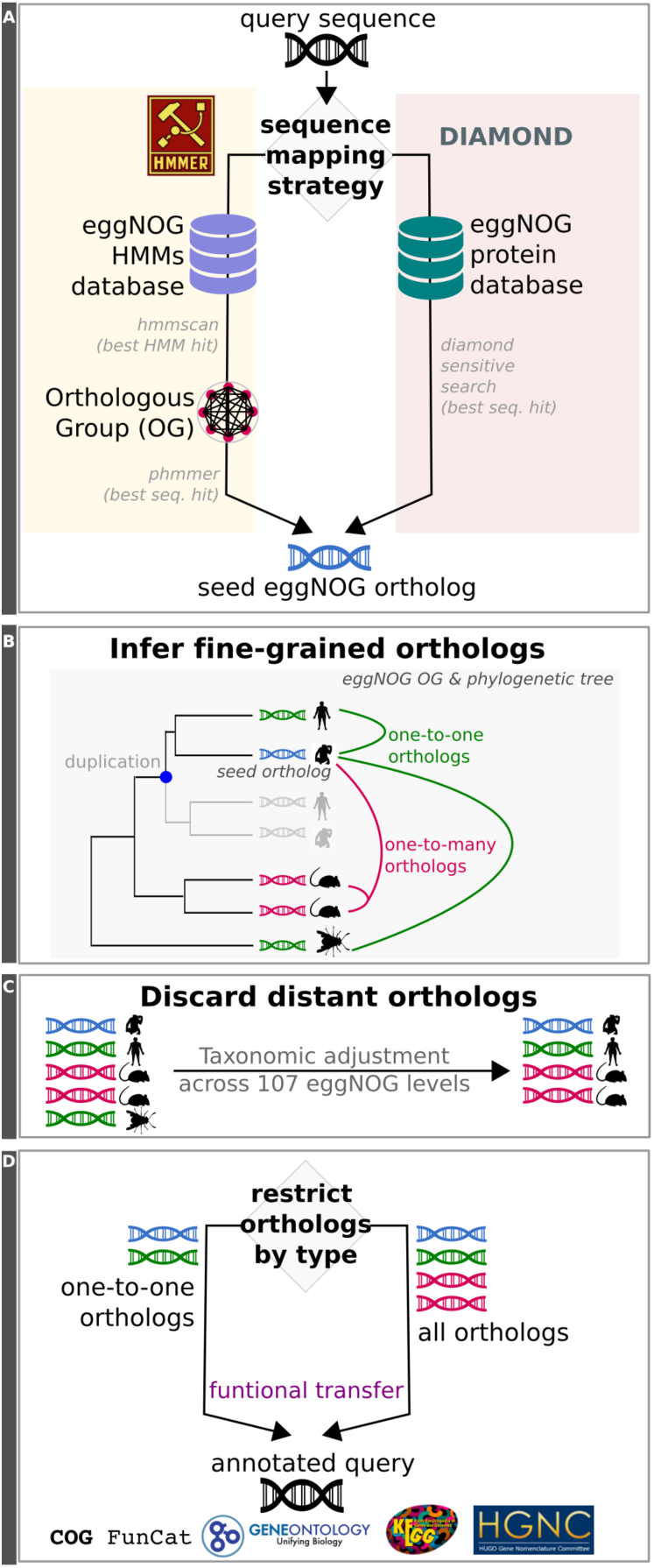
eggNOG-mapper workflow. Schematic representation of the eggNOG-mapper workflow and its different execution modes. **A)** Sequence mapping step showing two available options: HMM-based searches (left), and DIAMOND-based searches (right). For each query, both options lead to the best seed ortholog in eggNOG. **B)** Inference of fine-grained orthologs based on the precomputed eggNOG phylogenies associated to the Orthologous Groups (OG) where the seed orthologs are found. **C)** Fine grained orthologs are further filtered based on taxonomic criteria. Distant orthologsareautomatically excluded unless manually specified **.D)** Functional transfer is performed using either one-to-one orthologs or all available orthologs. Gene Ontology terms, KEGG pathways, COG functional categories and predicted gene names are transferred from orthologs to query.

## Accuracy of functional assignments

To test the performance of annotation transfer along orthology relationships, we benchmarked eggNOG-mapper GO predictions for the complete proteomes of five functionally well-characterised model organisms alongside those produced by two existing approaches. The first is standard BLAST homology searches at different E-value thresholds, which is the approach used by tools like Blast2GO (Götz et al. 2008) and RAST (Overbeek et al. 2014). The other is the state-of-the-art InterProScan 5 pipeline (Jones et al. 2014), which unifies twelve independent databases into a manually curated collection of functional models based on sequence profiles.

As a gold standard for functional assignment, we used curated GO terms (evidence code other than “IEA” or “ND”) as true positives, and curated taxon exclusion GO data (Deegan née Clark et al. 2010) as false positive terms. GO terms not falling into the true or false positive categories were considereduncertainassignments. Specieswereselectedonbasisofsufficientexperimental annotation deposited in public databases, thus ensuring a high coverage of curated GO terms per protein. BLAST-based annotation was performed using the same set of reference proteomes and functional data as in eggNOG v4.5. In addition, we excluded each target proteomefromall reference databases when annotating that species, both for eggNOG-mapper and BLAST, and disabled the automatic taxonomic adjustment in eggNOG-mapper to not unfairly penalize BLAST. This experimental setup allowed us to measure the specific effect of excluding paralogs from the functional transfer process. On the other hand, as we could not exclude self-annotations from InterProScan, the comparison with eggNOG-mapper was done using default parameters of both programs. This setup is notrepresentative ofmeasuring the absolute amountof curated terms recovered per protein, as circularity in annotations cannot be prevented without deep alterations to the InterProScan code or data. However, it allowed us to evaluate the rates of false and uncertain assignments achieved by eggNOG-mapper per protein, as compared with those based onthe manually curated InterProScan functional models (see Materials and Methods for more details).

Compared with BLAST-based annotations at the most stringent E-value cutoff tested (1E-40), eggNOG-mapper increased the proportion of true positives (term assignments validated through manual curation) to false positive term assignments per protein by 7% on average (Figure 2, left panel). Similarly, the proportion of curated terms recovered over total assignments(which also includes terms neither supported by curation nor excluded by taxonomy) improved by 19% using eggNOG-mapper (Figure 2, middle panel). BLAST-based annotation covered a larger portion of the target proteomes (Figure 2, right panel), but at the cost of considerably lower quality of annotations compared to eggNOG-mapper: i.e. the latter annotated 9% more proteins with only true positives assignments (Figure 2, blue bars in right panel) and 2% fewer proteins with only false or uncertain assignments (Figure 2, orange bars in right panel). These results were consistently achieved regardless of the target species or E-value threshold (applied to both BLAST and eggNOG-mapper hits to achieve a fair comparison). However, we found even more marked differences at lower E-value cutoffs (0.001 and 1E-10 cutoff bars in Figure 2).

**FIGURE 2.**
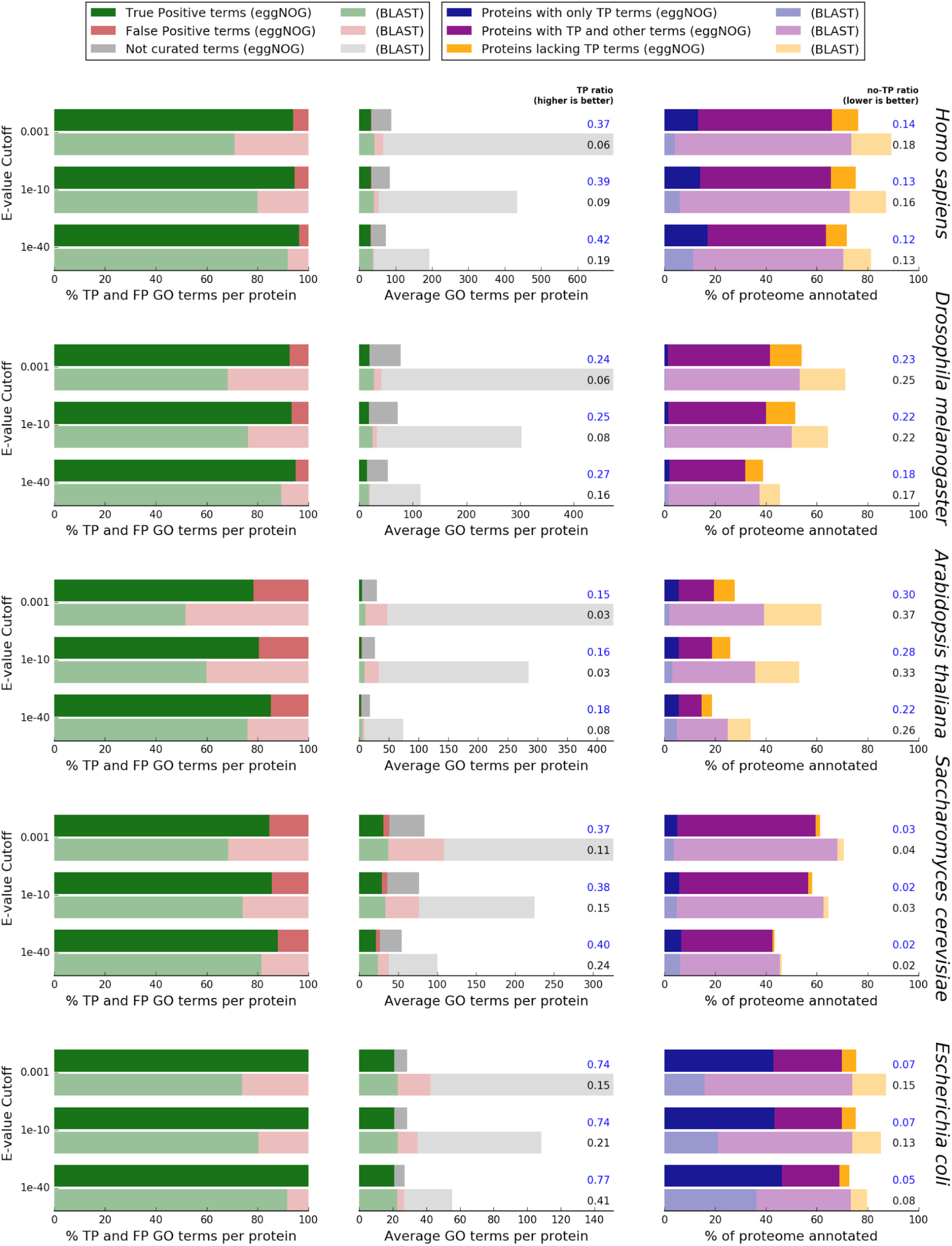
BLAST vs eggNOG-mapper. Comparison of the annotation results for five model species using eggNOG-mapper(brighter colours) and BLAST (dimmed colours). **Left panel** shows the per-protein average proportion of true positive GO term assignments (TP, green, supported by manual curation) to false positive term assignments (FP, red, derived fromtaxonomicexclusioncriteria).Withineachplot, consecutive pairs of horizontal bars represent different BLAST E-value cutoffs ranging from 1E-03 to 1E-40, with sequence matches under this cutoff being excluded from both BLAST and eggNOG-mapper hits. **Middle panel** shows the per-protein average number of true positive GO term assignments (green), false positive term assignments (red), and assignments of GO terms where neither curated evidence nor taxonomic exclusion criteria holds (grey). Next to the plot is shown the ratio of true positive term assignments over the total number of assignments (TP ratio). **Right panel** shows the percentage of each proteome that receives annotation, indicating the fraction of proteins that were annotated exclusively with curated true positive terms (TP, blue); proteins annotated with curated terms but also false or uncertain assignments (purple); and proteins that only received false or uncertain assignments (no-TP, orange).

Finally, webenchmarkedeggNOG-mapper annotations against those producedby InterProScan v5.19-58 (Jones et al. 2014). As the sources for GO annotations could not be adjusted in InterProScan, circularity in the annotation of reference proteomes could not be avoided. Therefore we compared the performance of both tools without any additional cutoff and using default parameters. On average, the total number of terms assigned by eggNOG-mapper per protein was 32 times higher than with InterProScan. These annotations were inferred with a similar ratio of false positive assignments (0.3% difference, Figure 3, left panel). In addition, the ratio of true positive terms over total assignments was increased by 26% in eggNOG-mapper (Figure 3, middle panel). Except for *Arabidopsis thaliana,* proteome coverage was also improved. eggNOG-mapper rendered 30% more proteins receiving only true assignments (Figure 3, blue bars in right panel), and 32% less proteins having false or uncertain terms only (Figure 3, orange bars in right panel).

**FIGURE 3.**
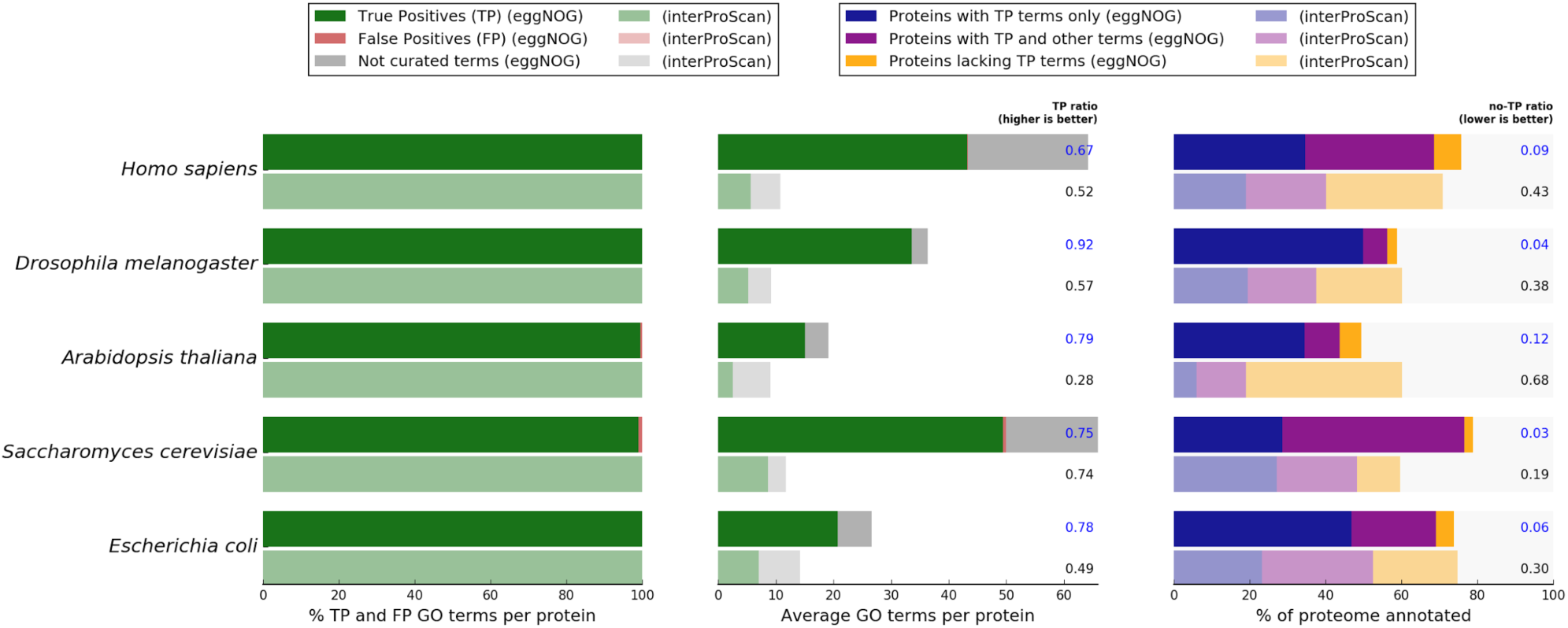
InterProScan vs eggNOG-mapper. Comparison of the annotation results for five model species using eggNOG-mapper (brighter colours) and InterProScan (dimmed colours) with default parameters and without further restrictions. The **left panel** shows the per-protein average proportion of true positive GO term assignments (TP, green, supported by manual curation) to false positive term assignments (FP, red, derived from taxonomic exclusion criteria). Consecutive pairs of horizontal bars represent each species in the benchmark. The **middle panel** shows the per-protein average number of true positive GO term assignments (green), false positive term assignments (red), and assignments of GO terms where neither curated evidence nor taxonomic exclusion criteria hold (grey). Next to the plot is shown the ratio of true positive term assignments over the total number of assignments (TPratio).The **right panel** shows the percentage of each proteome that receives annotation, indicating the fraction of proteins that were annotated exclusively with curated true positive terms (TP, blue); proteins annotated with curated terms but also false or uncertain assignments (purple); and proteins that only received false or uncertain assignments (no-TP, orange).

Computation time was also considerably reduced. Overall, eggNOG-mapper completed annotations ~15 times faster than running BLAST and 2.5 faster than InterProScan using the same system and the same number of CPU cores. In the context of this benchmark we disabled the lookup service in InterProScan, since it would not improve the speed when annotating novel proteomes.

## Conclusions

Although orthology is considered one of the most reliable sources for functional transfer, computational requirements, as well as the lack of practical tools, have hindered its use for the functional annotation of novel genomes. Here, we have presented a novel method and a tool, eggNOG-mapper, for easily annotating large sets of proteins based on fast orthology mappings.

We observed clear improvements relative tohomology-basedannotationsusingBLAST, reinforcing the central idea of orthologs being better functional predictors than paralogs, as well as showing how the latter cannot be fully excluded merely by using strict E-value BLAST thresholds. On the other hand, eggNOG-mapper achieved the same low rate of false positives as when using manually curated InterProScan functional models, while still increasing the amount and quality of annotations. Furthermore, eggNOG-mapper runs orders of magnitude faster than a standard BLAST-based approach, and at least 2.5 faster than InterProScan, whichmakesitparticularly suitable for large-scale annotation projects such as in metagenomics.

eggNOG-mapper is distributed as a standalone package and can be easily integrated into third-party bioinformatics pipelines. In addition, we provide an online service that facilitates functional annotation of novel sequences by casual users (http://eggnog-mapper.embl.de). The tool is synchronised with the eggNOG database, ensuring that the annotationsourcesandtaxonomic ranges will be kept up-to-date with future eggNOG versions.

## Material and Methods

### Benchmark data

Benchmarking was performed using the proteomes of five model species downloaded from eggNOG v.4.5, namely *Escherichia coli, Drosophila melanogaster, Saccharomyces cerevisiae, Arabidopsis thaliana* and *Homo sapiens*. For all five proteomes, GO terms were retrieved from eggNOG version 4.5. GO terms with evidence codes different than IEA or ND were considered curated positive terms. Similarly, any assignment of a term to a protein from a taxon it is excluded from (according to taxon exclusion data downloaded in Dec 2015 from the Gene Ontology Consortium) was considered a false positive (e.g. nervous system development terms assigned to a plant gene). Non-curated terms that are not explicitly listed in the false positive category were considered uncertain terms.

### Benchmark setup: BLAST

BLAST searches were performed using NCBI-BLAST 2.3.0 with an E-value threshold of 0.001, 20 threads and unlimited number of hits. eggNOG v4.5 was used as target database (http://eggnogdb.embl.de/download/eggnog_4.5/eggnog4.proteins.core_periphery.fa.gz). While annotating query sequences, self hits were excluded both from BLAST hits and eggNOG-mapper hits to avoid circular annotations. No taxonomic restrictions were applied when transferring Gene Ontology (GO) terms from BLAST or eggNOG-mapper hits (automatic taxonomic adjustment was manually disabled).

### Benchmark setup: InterProScan

InterProScan-5.19-58.0 (Jones et al. 2014) was run for all reference proteomes with default options and enabling GO annotation: *“--goterms --iprlookup -pa”*. All GO term predictions from all InterProScan source categories were used. eggNOG-mapper was executed with default options, which include using all types of orthologs and automatic adjustment of taxonomic sources. To standardize results and make the two set of predictions fully comparable, each GO term obtained from either program was augmented to include all its parent GO terms in theGeneOntology hierarchy. For speed comparisons, both programs were executedenablingtheuseof20CPU cores.

## SupplementaryMaterial

For reproducibility, benchmark scripts and raw data are provided as online supplementary material at http://github.com/jhcepas/emapper-benchmark

